# BARtab & bartools: an integrated Nextflow pipeline and R package for the analysis of synthetic cellular barcodes in the genome and transcriptome

**DOI:** 10.1101/2023.11.21.568179

**Authors:** Henrietta Holze, Laure Talarmain, Katie A. Fennell, Enid Y. Lam, Mark A. Dawson, Dane Vassiliadis

## Abstract

Cellular barcoding using heritable synthetic barcodes coupled to high throughput sequencing is a powerful technique for the accurate tracing of clonal lineages in a wide variety of biological contexts. Recent studies have integrated cellular barcoding with a single-cell transcriptomics readout, extending the capabilities of these lineage tracing methods to the single-cell level. However there remains a lack of scalable and standardised open-source tools to pre-process and visualise both population-level and single-cell level cellular barcoding datasets. To address these limitations, we developed *BARtab*, a portable and scalable Nextflow pipeline that automates upstream barcode extraction, quality control, filtering and enumeration from high throughput sequencing data; and *bartools*, an open-source R package that streamlines the analysis and visualisation of population and single-cell level cellular barcoding datasets. *BARtab* contains additional methods for the extraction and annotation of transcribed barcodes from single-cell RNA-seq and spatial transcriptomics experiments, thus extending this analytical toolbox to also support novel expressed cellular barcoding methodologies. We showcase the integrated *BARtab* and *bartools* workflow through comparison with previously published toolsets and via the analysis of exemplar bulk, single-cell, and spatial transcriptomics cellular barcoding datasets.

## Main

Modern cellular barcoding approaches can accurately trace clonal lineages by labelling individual cells within a population with unique and heritable genetic barcodes that are subsequently read out using high throughput sequencing technologies ^1–3^. These techniques enable the investigation of clonal dynamics at unprecedented scale, helping to map developmental trajectories and lineage relationships across multiple organisms and experimental systems ^2–5^. A fundamental principle of cellular barcodes is that they are heritable through cell division such that each daughter cell inherits the same barcode as its parent, thereby establishing a clonal lineage. Typically, this is achieved by engineering a unique barcode into the genome of each cell. Cellular barcoding techniques most commonly employ viral vectors or recombinant transposases to introduce a complex library of synthetic barcode sequences into the genomes of a target cell population, resulting in the unique labelling of hundreds to thousands of individual cells ^6^. For population-level lineage tracing studies, the resulting barcodes can be isolated from genomic DNA by PCR, sequenced using a high throughput sequencing platform and enumerated to reveal the frequency of each clone in a population. With the advent of single-cell sequencing technologies, new methods have incorporated the readout of synthetic barcodes into single-cell transcriptomic datasets ^7–13^. Here, synthetic barcodes are cloned into a reporter gene cassette such that they are present on mature mRNA transcripts and can be read out using polyA capture based single cell RNA-seq protocols. This concept also extends to sequencing-based spatial transcriptomics technologies that employ a similar polyA based mRNA capture strategy to link gene expression with spatial context *in situ* ^14^.

While cellular barcoding is a powerful methodology for understanding clonal dynamics across time and space, the analysis of barcoding datasets can be complex, causing many to turn to bespoke data analysis tools. Despite the maturity of the cellular barcoding field, there remains no accepted gold standard data analysis pipeline or workflow suitable for population-level datasets, let alone cellular barcoding analysis from single-cell or spatial datasets ^15^. Recent efforts to standardise the analysis of such data have focussed primarily on dataset visualisation and lack support for the upstream pre-processing of raw data or for the next wave of lineage tracing studies that will utilise expressed synthetic barcodes ^16–18^. Indeed, recent single cell expressed barcoding studies incorporate their own customised analysis pipelines in a manner that lacks versatility for studies that utilise a conceptually similar yet slightly different biological workflow ^7,8,10,11,19^. Thus, there is a need for an end-to-end integrated solution for cellular barcoding dataset pre-processing, analysis and visualisation that is flexible to different barcode designs and is portable and scalable across computational environments.

To support the standardisation of barcode dataset pre-processing and quality control, here we introduce *BARtab*, a portable and scalable Nextflow pipeline which allows the generation of barcode counts tables from population level barcode sequencing workflows as well as barcode extraction, enumeration, and cell annotation from single-cell and spatial transcriptomics datasets. Moreover, to facilitate the downstream analysis and visualisation of cellular barcoding workflows at bulk and single-cell resolution we developed *bartools*, a flexible open-source R package that incorporates workflows for population-level cellular barcoding data analysis, as well as methods for single-cell expressed barcode analysis and visualisation. *bartools* provides a convenient interface between cutting-edge methods for the analysis of cellular barcoding datasets, and the robust analytical framework established within the R ecosystem. We demonstrate the improved performance and versatility of *BARtab* compared to other barcode pre-processing software and showcase the capabilities of our integrated *BARtab* and *bartools* workflow through the analysis and visualisation of exemplar population-level, single-cell level and spatial transcriptomics based cellular barcoding datasets which we make publicly available. Together, *BARtab* and *bartools* comprise an end-to-end integrated toolkit that will help streamline and standardise cellular barcoding experiments for the lineage tracing field at large.

## Results

### BARtab and bartools comprise an integrated cellular barcoding analysis workflow

Population-and single-cell-level cellular barcoding workflows are usually readout using high throughput sequencing, resulting in raw sequence data containing barcode information that must undergo quality control, barcode extraction, and quantification. This task is usually performed by bespoke software specific to the study in question. Publicly available tools that exist, such as genBaRcode ^16^, pycashier ^20,21^ or xcalibr ^22^, are limited in their flexibility for different barcode designs and lengths, support for paired-end datasets, portability and resource allocation, support for reference-based barcode quantification or support for single cell and spatial transcriptomics expressed cellular barcoding datasets (**Table 1**). In contrast, *BARtab* is an open-source Nextflow pipeline ^23^, written as a versatile, portable, reproducible, and scalable solution for high throughput barcode dataset pre-processing from population-level, single-cell and spatial transcriptomics cellular barcoding experiments. *BARtab* leverages widely used bioinformatics tools including *fastx-toolkit*, *FLASh*, *cutadapt*, *samtools*, *bowtie*, *starcode*, *umi-tools*, and *FastQC* ^24–30^. In its simplest form, running on population-level cellular barcoding data, *BARtab* performs the following steps: 1) Import and quality control of raw sequence data, 2) Barcode quality control and filtering, 3) Adapter trimming and extraction of barcodes from raw sequencing reads, 4) barcode quantification, and 5) Reporting (**Figure 1**).

**Figure 1.**
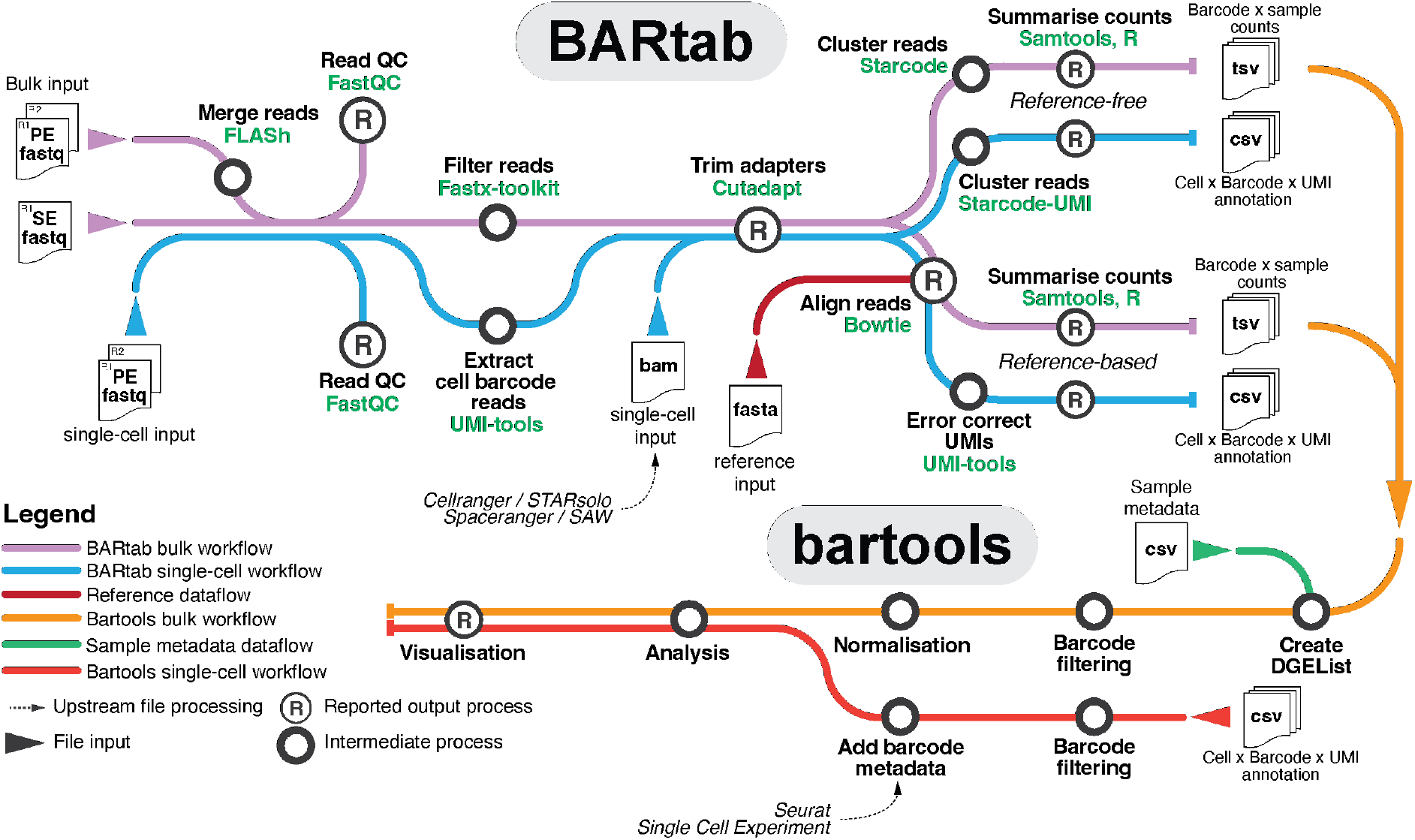
The BARtab & bartools workflow for high throughput cellular barcoding analysis. BARtab is a Nextflow pipeline to process high throughput sequencing datasets containing cellular barcode information. BARtab contains two main sub-workflows that permit the analysis of (1) single-end or paired-end population-based cellular barcoding data (“bulk workflow”), and (2) paired end single-cell datasets (“single-cell workflow”). For the bulk workflow, BARtab takes single or paired-end datasets in fastq format as input and performs read merging (paired-end only) quality filtering and adapter trimming (single and paired-end) and barcode quantification. Quantification of barcodes can occur using a reference-based (recommended) strategy which aligns putative barcode sequences to user supplied reference of known lineage barcodes, or by a reference-free workflow which utilizes a clustering-based approach within Starcode to merge similar sequences together based on a defined Levenshtein distance threshold. For the single-cell workflow, raw paired-end datasets in fastq format (e.g. from targeted amplification of cellular barcodes within a single-cell library) or aligned reads in BAM format (output as part of the pre-processing of single-cell data) are taken as input, barcode containing reads are selected and quality filtered and finally quantified using either a reference-based as above or via a reference - free approach using Starcode-UMI. Outputs of all workflows include tables of processed counts with individual barcodes as rows and sample counts as columns, as well as a MultiQC run report including read filtering and alignment information. Tables of counts from BARtab, alongside associated sample metadata, can be imported into bartools, for further downstream analysis and visualisation. For single-cell datasets, barcode import interfaces directly with popular R-based single-cell frameworks including Seurat and SingleCellExperiment resulting in cell level annotation with lineage barcode metadata which can be leveraged for analysis and visualisation in bartools and other single cell analysis packages.

**Table 1.**
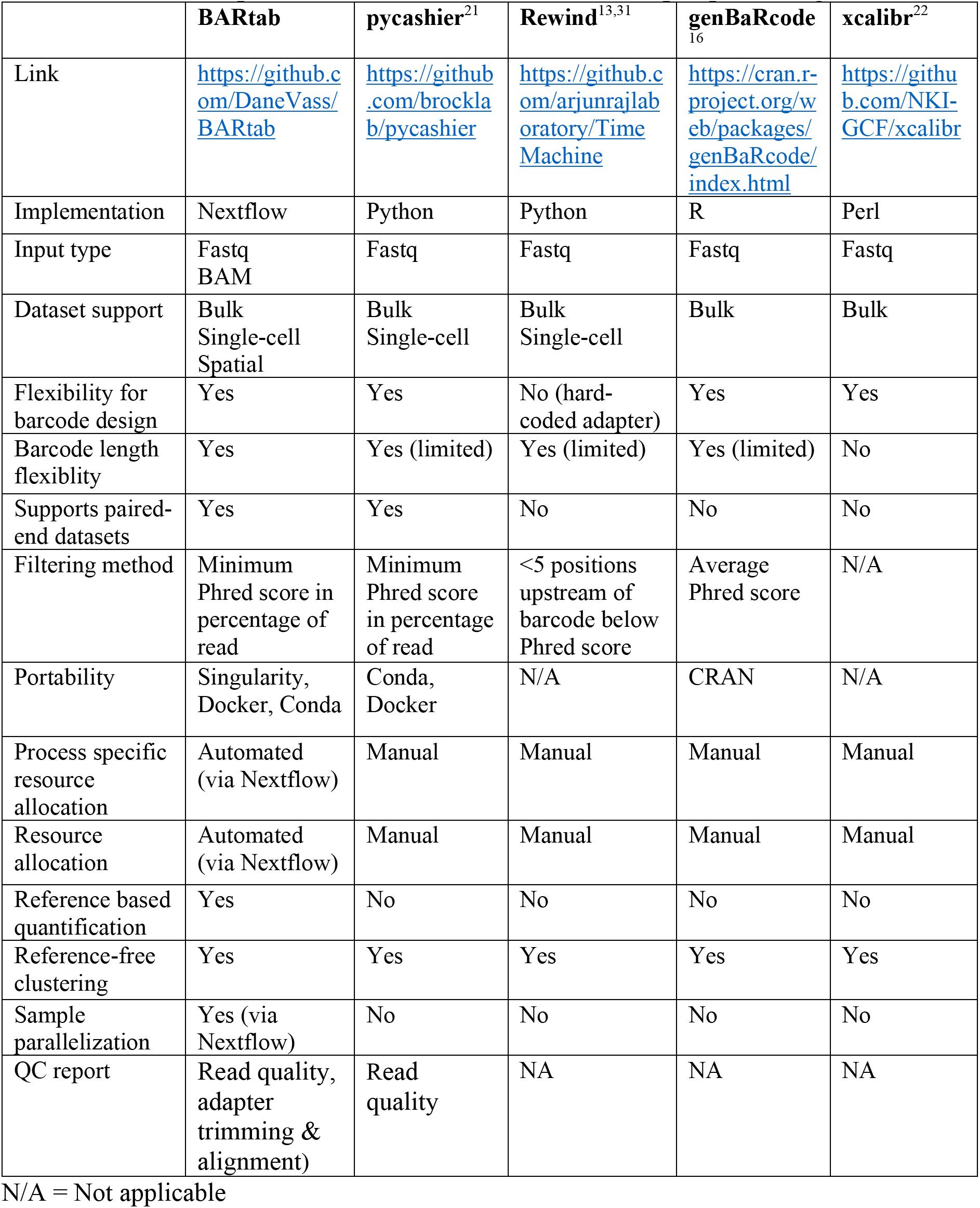
Feature comparison of BARtab and other barcode pre-processing tools.

Alternate sub-workflows are also available for situations where no reference library of barcodes is available, for paired-end reads that require merging prior to extraction (e.g. for barcode constructs where the length of the barcode is greater than the length of the sequenced read), or for extraction of barcode reads from single-cell and spatial transcriptomics datasets utilising expressed barcode technology (**Figure S1A**). For population-level datasets, the primary output of *BARtab* is a table of raw counts per barcode, where rows are individual barcodes and columns are individual samples. For single-cell datasets, *BARtab* outputs a single table per sample containing unique molecular identifier (UMI) and lineage barcode information per cell ID. This table can be imported as sample metadata into established R or Python-based single cell RNA-sequencing analysis packages such as *SingleCellExperiment* from the Bioconductor project, *Seurat* or *Scanpy* ^32–34^. For bulk-level data, the counts table output from *BARtab* is preformatted for, and can be easily read into *bartools,* thus connecting dataset pre-processing to downstream analysis and visualisation.

*Bartools* is an open-source R package that accepts tables of raw counts (with individual barcodes / tags as rows and samples as columns), such as those generated by *BARtab*, or other software as input. Utilising the *edgeR* DGEList framework ^35^, barcode count datasets can be read into *bartools* as individual count table files or as defined in an experimental samplesheet which organises raw count data alongside associated sample metadata (see **Table S1** for an example). Quality control metrics including the results of filtering and reference mapping stages from an associated *BARtab* pre-processing run can be plotted per sample in *bartools* using the *plotBARtabFilterQC()* and *plotBARtabMapQC()* functions. Following initial dataset filtering and QC, cellular barcoding datasets can be further processed using built-in functions that perform sample normalisation, downstream abundance & diversity analysis, and visualisation of synthetic barcode data (**Figure 1**). The *bartools* package also includes analysis methods that accept single-cell objects containing cellular barcoding information in Seurat or SingleCellExperiment format which can be used to aid in quality control and assess clone level properties in concert with popular scRNA-seq analysis workflows (**Table 2**). Overall, the integrated combination of *BARtab* and *bartools* improves upon other available dataset pre-processing and analysis / visualisation software by providing an end-to-end analysis solution that extends on previous offerings and permits analysis of single-cell and spatial expressed cellular barcoding datasets.

**Table 2.** Feature comparison of bartools and other barcode analysis toolkits.

|  | **bartools** | **genBaRcode** | **barcodetrackR** | **CellDestiny** |
| --- | --- | --- | --- | --- |
| Data structure | DGEList (edgeR) | custom R object | SummarisedExperiment | Rshiny |
| Normalisation modes | Percentage abundance, CPM, TMM | NA | Percentage abundance, CPM | Percentage abundance |
| QC and aggregation of replicates | Yes | No | No | Yes |
| Clone size analysis | Yes | No | Yes | Yes |
| Support for single-cell experiments | Yes | No | No | Yes |
| Rshiny app | No | Yes | Yes | Yes |

### BARtab improves upon previous cellular barcode dataset pre-processing software

Cellular barcoding datasets are prone to technical artifacts that skew the representation and perceived abundance of barcoded clones of interest. Barcode amplification by PCR, high-throughput sequencing, read filtering, and downstream analysis stages all present potential sources of technical error in a cellular barcoding workflow ^36,37^. In *BARtab*, we encourage a reference-based approach for the quantification of cellular barcoding datasets by mapping putative barcode containing reads to a known reference set of accepted barcodes ^1,2,9^. This reference set of barcodes can be obtained via deep sequencing of the library plasmid pool, enabling comprehensive identification of true barcodes present within a particular library without overt reliance on sequence error correction processes. In contrast, similar approaches including genBaRcode ^16^, pycashier ^20,21^ or xcalibr (https://github.com/NKI-GCF/xcalibr) take a distance threshold approach to combine similar barcodes together that can arise through errors during PCR or sequencing ^16,38^. Although these approaches sidestep the requirement of a deeply sequenced reference, further quality control and filtering procedures are recommended to eliminate spurious results. Alongside the reference-based workflow, *BARtab* can also perform reference-free identification and quantification of barcodes from single-cell and bulk datasets by employing a Levenshtein distance-based clustering approach implemented in Starcode ^24^. In summary, *BARtab* allows for reference-based as well as reference-free extraction of barcodes from both bulk and single-cell datasets, is flexible towards barcode design and sequencing quality, provides a detailed quality report and is portable and parallelized to facilitate large-scale data processing.

To demonstrate the versatility of *BARtab*, we compared its performance to pycashier and Rewind/TimeMachine, two recently published toolkits for cellular barcode extraction ^13,21,31^. Using each tool, we re-analysed an exemplar population-level cellular barcoding dataset consisting of 22 individual samples from a recent Rewind/TimeMachine study by Goyal et al.^31^. Since the pycashier and Rewind/TimeMachine approaches do not support reference-based barcode quantification (**Table 1**), we ran *BARtab* using the reference-free clustering approach. We used default parameters for the Rewind/TimeMachine tool as per the original publication and parameters for *pycashier* and *BARtab* that matched those analysis conditions as closely as possible to allow comparable barcode quantification performance.

We observed a total of 144745 barcodes (97.3%) with at least 0.001% frequency within a sample detected by all three tools across the 22 samples. A further 1810 barcodes (1.2%) were only detected by TimeMachine and BARtab. In contrast, just 92 barcodes (0.1%) were only detected by the original TimeMachine analysis and pycashier (**Figure S1A**). The improved detection rate of BARtab is likely due to its increased flexibility for variable barcode lengths (see Methods). We also examined barcode quantification concordance of BARtab to the published dataset. Here we observed striking concordance in barcode representation and abundance with Pearson and Spearman correlation values between the two tools greater than 0.96 for all samples (**Figure S1B**). Finally, we compared the runtime of BARtab, pycashier and Rewind/TimeMachine across the 22 sample dataset on a computing cluster, with 32 GB of memory and 20 CPUs allocated to each workflow. Here, BARtab and pycashier performed comparably, processing all datasets in 40 and 36 minutes respectively, whilst Rewind/TimeMachine took 4 hours 59 minutes. The stark difference in runtime can be attributed to the lack of parallelization and resource allocation support in TimeMachine causing all samples to be run serially. Since BARtab produces comprehensive QC reports detailing the results of raw data quality filtering and barcode extraction stages, we believe that the minimal increase in runtime compared to pycashier is offset by the additional benefit of this improved reporting. Together, these analyses highlight the versatility of BARtab for different cellular barcoding methodologies, and its improved sensitivity and processing speed compared to currently available software.

### Bartools supports the quality control of population-level cellular barcoding datasets

Next, to demonstrate the capabilities of *BARtab* and *bartools* for both bulk and single-cell level cellular barcoding analyses, we generated a population-level cellular barcoding dataset using the SPLINTR lineage tracing system ^9,39^. Here, Acute Myeloid Leukaemia (AML) cells were cultured in the presence of gradually escalating doses or an upfront high dose of Cytarabine (AraC), a conventional chemotherapy used routinely in the clinic, or IBET-151, a targeted epigenetic therapy against the BET bromodomain family of transcriptional co-activators which has shown pre-clinical efficacy against several AML subtypes ^39^ (**Figure 2A**). This “dose-escalation” dataset comprises bulk barcode-seq data per dose and timepoint.

**Figure 2.**
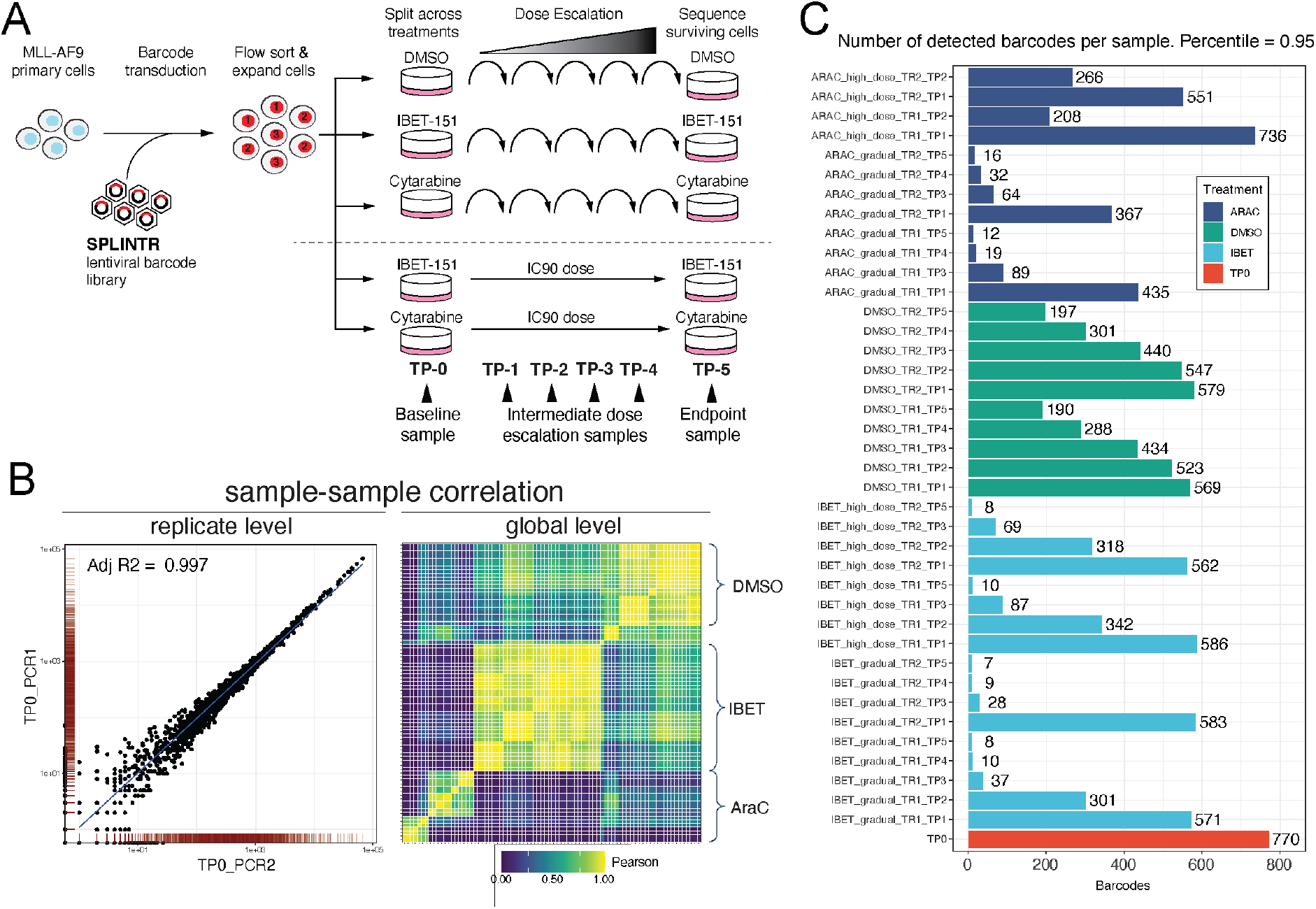
Cellular barcoding quality control analysis of the dose response dataset. **A)** Schematic of dose escalation dataset experimental design. Drug naïve MLL-AF9 cells were transduced with SPLINTR barcode encoding lentivirus in liquid culture, sorted for barcode expression and expanded for one week in liquid culture. Following expansion, barcoded clones were split evenly into media containing either 400nM IBET-151, 300nM AraC or 0.1% (v/v) DMSO (vehicle control) such that each treatment arm received an identical barcode repertoire. Cells were re-plated weekly in escalating concentrations of IBET-151/AraC or vehicle. Simultaneous to the dose escalation, the same pool of cells were seeded into an IC90 dose of IBET151 (800nM) or AraC (700nM). Barcode sequencing was performed on samples obtained at the timepoints indicated. The experiment was performed in biological duplicate. **B)** Replicate and global level sample correlation analyses. Left; Representative scatterplot of barcodes in technical replicates for the TP0 sample. Right; pairwise Pearson correlation heatmap of all biological replicate samples (indicated) in the dose escalation dataset. **C)** Total number of barcodes present at the 95th percentile in each sample following sample QC and merging of technical replicates.

Following the pre-processing of raw reads into barcode counts and their import into *bartools*, samples can be filtered according to absolute and relative (proportion) thresholds. Total read depth and number of detected barcodes can then be assessed per sample. To demonstrate this, we applied both sample level and barcode level filters the dose-escalation dataset. First, we filtered samples using a 5^th^ percentile outlier threshold calculated from the total read counts across samples. This eliminated six low quality samples that were unlikely to yield reliable data (**Figure S2A**). Next, using the *thresholdCounts()* function, we assessed different relative and absolute read count thresholds on a per barcode basis and chose to apply a low-stringency filter, removing barcodes from the dataset with less than 10 reads across at least four samples. This eliminated 110 low abundance barcodes from the dataset that are likely the result of sequencing errors or other technical noise. This filtering resulted in a dataset of 1680 high-confidence barcodes across 41 samples. We observed a gradual decrease in total barcode numbers across the IBET-151 and AraC dose escalation timecourse (**Figure S2B**), consistent with our earlier findings in this model system ^39^.

If the experimental design incorporates technical replicates, which we also encourage, further QC can be performed using *calcReplicateCorr()* to assess replicate correlation and removing samples with correlation scores below a reasonable threshold (**Figure 2B**). Poor technical replicate correlation scores can indicate sampling bias during sample preparation or sequencing coverage issues that may complicate dataset interpretation. Following sample QC and filtering, technical replicates displaying greater than 90% correlation at the sample level were averaged using *collapseReplicates()* and the number of barcodes comprising the 95^th^ percentile in each remaining sample was calculated using the *calcPercentileBarcodes()* and *plotDetectedBarcodes()* functions (**Figure 2C**). Finally, dataset normalisation was performed using a counts per million (CPM) transformation via *normaliseCounts()*. Trimmed mean of M-values (TMM) ^40^ and percentage abundance (PCT) based metrics are also available as normalisation options within *bartools*. Applied correctly, these quality control and normalisation approaches result in a clean cellular barcoding dataset that is ready for downstream analysis and visualisation.

### Bartools enables visualisation and comparison of global and individual clone abundance

Following dataset normalisation, global visualisation of cellular barcoding data is useful to understand the overall clonal composition of within and between samples. *Bartools* contains several visualisation options to assess global clonal repertoires. For example, bubbleplots (via *plotBarcodeBubble()* and *plotOrderedBubble()*, **Figure 3A**) give an overview of relative sample composition while informing on the proportional abundance of clones and the identities of overrepresented clones within samples. Heatmaps (via *plotBarcodeHeatmap()*), similar to those implemented in the barcodetrackR package ^17^ allow similar global overview and stratification of samples based on multiple metadata fields, however are simpler to interpret on a subset of the total dataset (**Figure 3B**). Timecourse studies such as the dose escalation dataset can additionally benefit from timeseries plots which can reveal the temporal nature of clonal fitness (*plotBarcodeTimeseries,* **Figure 3C**). Further to analyses of clone abundance, it is also of interest to calculate correlation or relatedness scores between samples in a barcoding dataset. *Bartools* leverages gold-standard linear methods to determine sample similarity including principal component analysis (PCA), sample-sample distance, and correlation methods (**Figure 3D** & **Figure 3E**). Moreover, *bartools* incorporates functions from the R packages *vegan* and *ineq* ^41,42^ to calculate various population diversity metrics including the Shannon, Simpson, Inverse Simpson, and Gini indices (**Figure 3F**).

**Figure 3.**
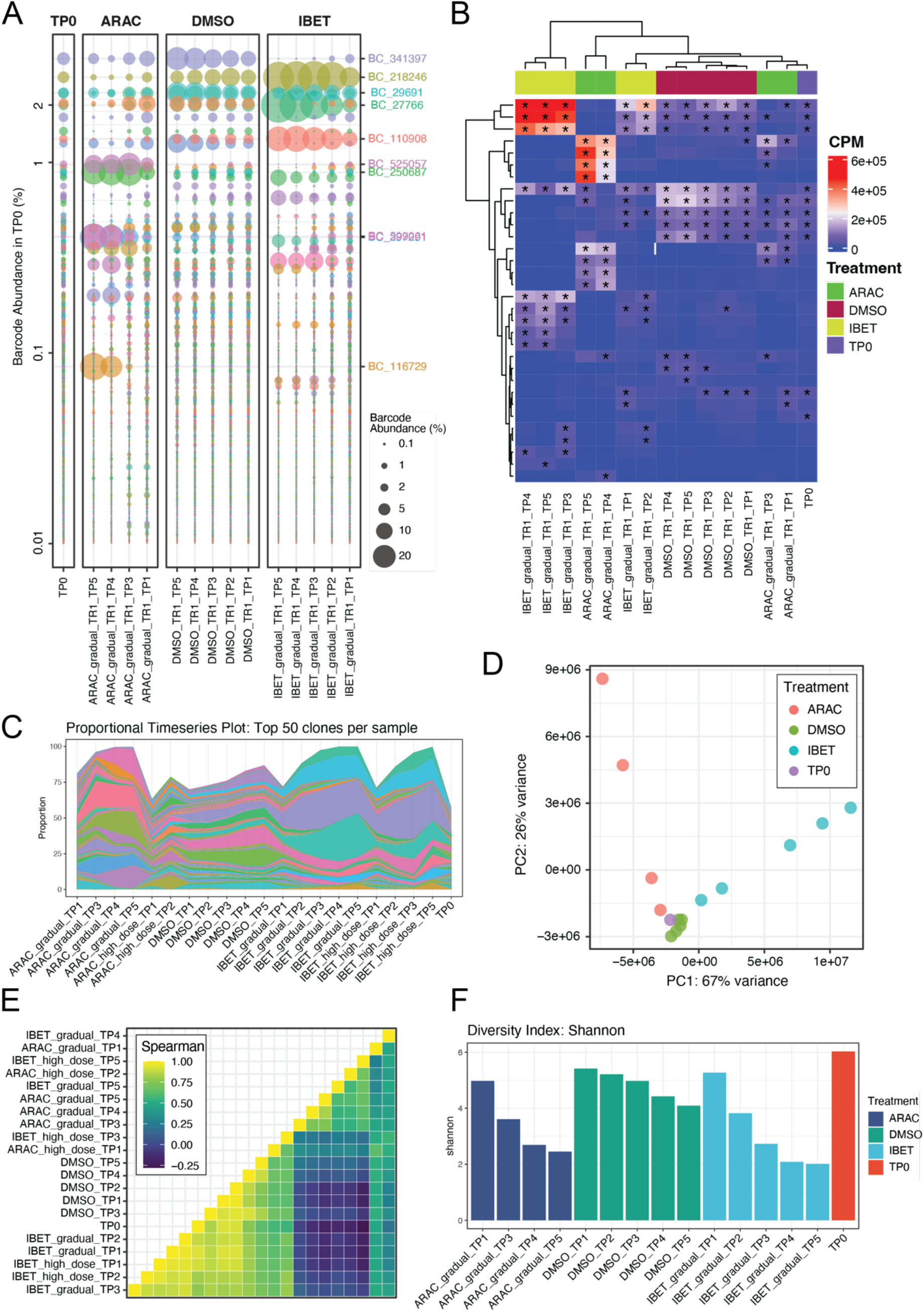
Analysis of global clonal repertoires, sample correlation and diversity in the dose escalation dataset. **A)** Bubbleplot of a subset of samples from the dose escalation dataset ordered in descending rank order according to percentage abundance in the baseline (TP0, red) sample. Left hand side y-axis indicates percentage abundance of each clone in the reference sample (TP0, red). Right hand y-axis indicates clones present at a percentage abundance above 5% in any sample. **B)** Heatmap of the dose escalation dataset showing counts per million (CPM) values for the top 10 most abundant barcodes (rows) in each sample (columns). Starred cells indicate barcodes that are among the top 10 most prevalent within that sample. **C)** Timeseries plot of the most abundant 50 barcodes across all samples within biological replicate 1 from the dose escalation dataset. **D)** Principal component analysis of the dose escalation dataset. Vehicle and drug (IBET-151 or ARAC) treatment conditions are clearly separated across PC1 while timepoint separates out on PC2. **E)** Sample-sample pairwise Spearman correlation matrix of the dose escalation dataset, hierarchically clustered according to sample similarity. **F)** Histograms of shannon diversity for baseline TP0, vehicle (DMSO) and drug treated (IBET-151 or ARAC) samples at each timepoint within biological replicate 1 from the dose escalation dataset.

To gain further insight into the clonal dynamics of drug resistance in the dose escalation dataset, we performed global analyses of the quality controlled and normalised dose escalation dataset using *bartools*. These analyses revealed clear outgrowth of distinct groups of clones in the vehicle and either drug treated conditions (**Figure 3A** & **B**). By PCA, treatments were clearly separated along principal component 1 (PC1) with the dose escalation timecourse separated along principal component 2 (PC2) (**Figure 3D**). These differences were also observed in correlation and diversity analyses across samples with later timepoints in the drug and vehicle conditions both less diverse and less well correlated with the baseline and early dose escalation samples (**Figure 3E** & **F**). These analyses reveal that IBET-151 or AraC therapies select for distinct groups of clones demonstrating the utility of bartools to extract biological insight from cellular barcoding datasets.

Following global level analyses of clonal composition within samples, it is of interest to interrogate the relative fitness of individual barcodes across samples or treatment groups. Such clone-level visualisations are also supported in *bartools* and can be useful to specifically compare clones of interest relative to others across samples or conditions. Using the *plotBarcodeBoxplot()* function, normalised abundance of individual clones can be displayed across sample metadata conditions. For the dose escalation dataset these analyses revealed striking differences in clonal fitness across the vehicle and treatment conditions with two classes of clones emerging: those that display increased fitness in vehicle conditions (**Figure S3A**), and those that display increased fitness in either drug treatment (**Figure S3B** & **S3C**). Overall, the combination of bulk and clone level analyses afforded in *bartools* will enable researchers to thoroughly interrogate clonal dynamics in diverse systems and under various perturbation conditions.

### BARtab and bartools facilitate single-cell expressed cellular barcoding data analysis

Recently developed expressed cellular barcoding tools, such as SPLINTR ^9^, Rewind/FateMap ^13,31^, Clonmapper ^21^, TREX ^14^, Watermelon ^10^, LARRY ^8^, and CellTag ^7,43^ utilise the 3’ capture of polyadenylated messenger RNAs in scRNA-seq libraries to capture transcripts encoding the cellular lineage barcode of each cell. Incorporation of these transcripts links cellular barcodes to a cell ID and unique molecular identifier (UMI) thus labelling single-cell transcriptomes with their corresponding clonal identity. We developed *BARtab* to be compatible with these expressed cellular barcoding workflows. The single-cell sub-workflow of *BARtab* accepts two types of inputs: 1) Sequence data in binary alignment map (BAM) format arising from single cell data pre-processed using the widely utilised CellRanger pipeline from 10X Genomics ^44^ or the open-source STARsolo ^45^; or 2) paired-end data in fastq format following targeted amplification of barcode containing reads from single-cell library cDNA (**Figure 1**).

Running in “single-cell” mode, *BARtab* first extracts reads containing lineage barcode information according to user defined parameters for barcode identification. For BAM file inputs, the cell ID and UMI are contained within the read tag information, allowing lineage barcode, cell ID and UMI information to be extracted simultaneously. For paired-end fastq inputs arising from targeted amplicon sequencing of the single cell library, cell ID and UMI information are extracted separately to the lineage barcode information and merged. Following read extraction, barcode containing reads are either mapped to a reference library of barcodes to annotate the clonal identity of a cell or processed using a reference-free approach similar to that employed in the population-level workflow (**Figure 1**). The pipeline outputs a cell metadata table containing cell ID, barcode, and number of UMI supporting the barcode annotation which can be imported into widely used R and Python based single-cell analysis packages including *Seurat*, *SingleCellExperiment* and *Scanpy*.

To demonstrate the single-cell annotation and analysis capabilities of *BARtab* and *bartools,* we applied the *BARtab* “single-cell” workflow to a 10X Genomics 3’ single-cell RNA-seq dataset consisting of 14086 SPLINTR barcoded murine AML cells cultured *in vitro* (hereafter the “single-cell” dataset) ^9^. Most cells identified with *BARtab* contained a single lineage barcode consistent with our previous results ^9^ (**Figure 4A**). Next, we imported our *BARtab* annotations into Seurat for further downstream analysis. Examining the raw dataset, we observed a positive relationship between total UMI counts or total detected features per cell and lineage barcode detection status suggesting that annotation of lineage barcodes could also assist with quality control of single cell datasets (**Figure 4B**). We reasoned that lineage barcode annotations could also be useful to diagnose biases related to barcode delivery or recovery in subpopulations of cells within a sample. We examined the percentage lineage barcode detection relative to the median UMIs detected per cluster. This analysis revealed no overt bias in the detection of barcode annotated cells across Louvain clusters by UMAP visualisation (**Figure 4C**). However, we did note that clusters with higher median overall transcript abundance in the single-cell dataset had an increased percentage of lineage barcode annotated cells supporting the positive relationship between total UMI counts and lineage barcode detection (**Figure 4D**).

**Figure 4.**
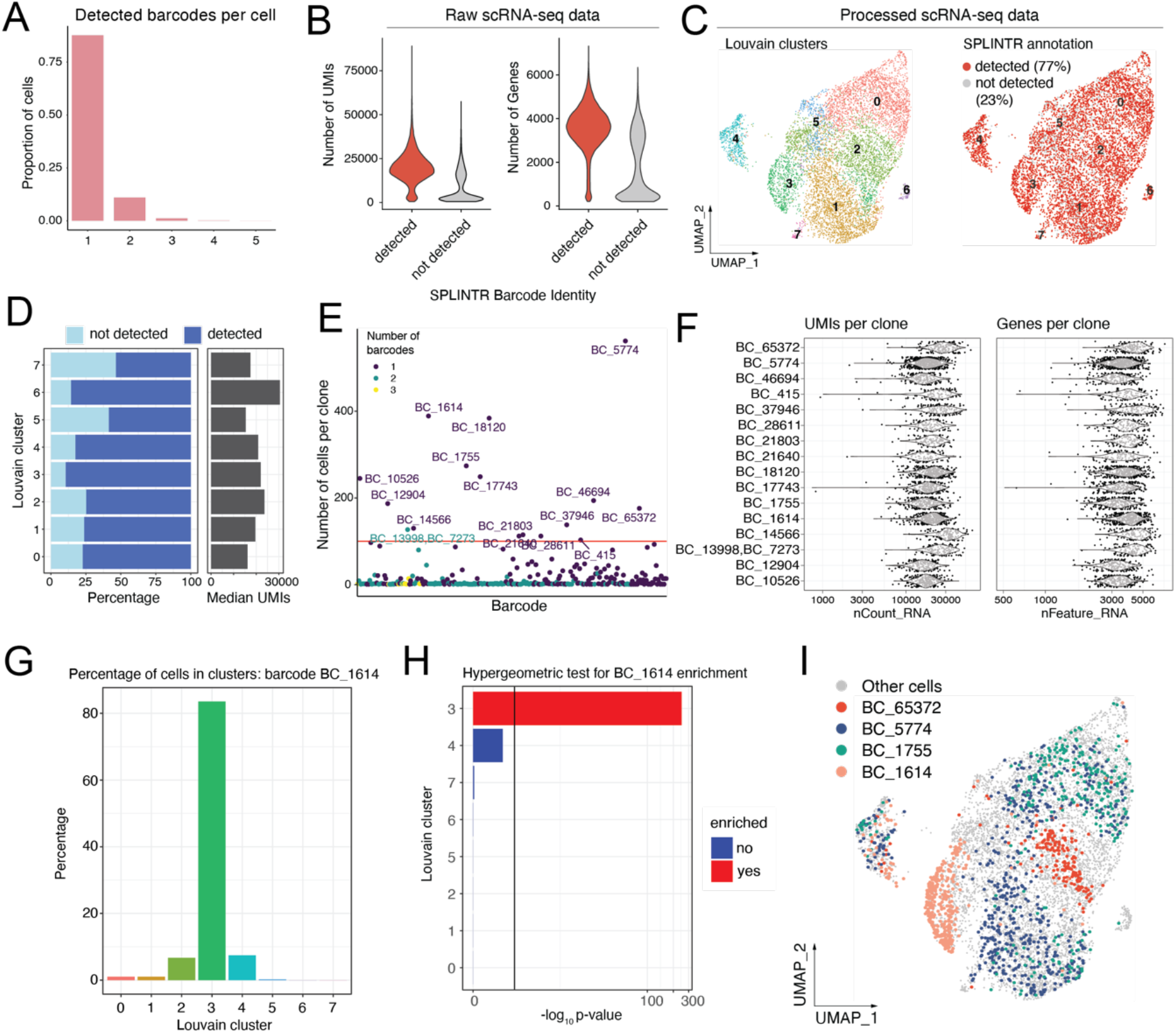
Expressed cellular barcode pre-processing and analysis with *BARtab & bartools*. **A)** Histogram showing detected barcodes per cell in the single-cell dataset. **B)** Number of UMIs detected per cell and number of genes detected per cell are shown for cells with a barcode detected (red) or not detected (grey). **C)** UMAP projections of the single-cell dataset following quality control, filtering and normalisation showing Louvain clusters (left) and expressed barcode detection status (right). **D)** (left) Percentage of cells with lineage barcode detected per 100 cells for each Louvain cluster. (right) Median UMIs per cell for each Louvain cluster. **E)** Dotplot showing the number of cells per lineage barcode. **F)** Violin plots of number of UMI counts per clone (nCount_RNA) and number of genes detected per clone (nFeature_RNA) for the 16 clones represented by 100 or more cells in the single cell dataset. **G)** Histogram showing the percentage of cells comprising an exemplar clone, BC_1614, within each Louvain cluster. **H)** Hypergeometric test results for enrichment of an exemplar clone, BC_1614, across Louvain clusters. **I)** UMAP visualisation of the single-cell dataset with cells from selected clones highlighted.

The inclusion of lineage barcode information in a single-cell dataset can reveal distinct properties of individual clones which can be related to phenotypic data. *Bartools* contains several functions to assess such clone level properties by leveraging widely used single-cell data structures within the R ecosystem (e.g. Seurat or SingleCellExperiment class objects). A simple yet powerful visualisation is the number of cells detected per barcode which we show for the single-cell dataset using the *plotCellsPerGroup()* function. For the single-cell dataset this analysis revealed 16 clones represented by at least 100 cells, some of which were characterised by multiple barcode integration events (**Figure 4E**). In addition, using the *plotMetrics()* function we could identify differences between clones for relevant metrics such as the number of individual transcripts (UMI’s) or features detected for each cell (**Figure 4F**). These analyses can help inform on functional or phenotypic differences between individual lineages.

The degree of transcriptional homo-or heterogeneity within clonal lineages is also of interest to many studies. *Bartools* incorporates percentage based (**Figure 4G**) and hypergeometric testing approaches at the cluster level via the *plotCellsInClusters() and plotClusterEnrichment()* functions to determine if certain lineages are enriched within different regions of transcriptional space. This analysis revealed distinct transcriptional patterns between the top 10 most represented clones, with some clones, such as BC_1614, localising primarily to a single Louvain cluster (**Figure 4H** & **4I**) suggesting transcriptional conservation in these clones despite an extended period of expansion in culture. Importantly, the *plotClusterEnrichment()* function is agnostic to the metadata variable and can be applied to any grouping of cells desired by the user. To demonstrate this, we analysed the enrichment of cells in different phases of the cell cycle per cluster. Cell cycle phase was annotated for the single-cell dataset using Seurat v4 and enrichment of cells in G1, G2M and S phase was analysed using *plotClusterEnrichment()* in bartools. This analysis revealed that the Louvain clusters overrepresented for actively growing/dividing cells (G2M/S phase) were also overrepresented for the most abundant clones (**Figure S4**). Overall, these quality control and analysis capabilities afforded by *bartools* can provide valuable insight into the nature of functional differences between different clonal lineages in a single-cell RNA-seq dataset as directed by genetic and/or non-genetic factors particular to biological system in question.

### Extending the cellular barcoding toolbox to spatial transcriptomics datasets

Recent advances in spatial genomics technologies have enabled the identification and sequencing of the endogenous spatial arrangement of individual cells *in situ.* 10X Genomics Visium and BGI Stereo-seq ^46^ are two recently developed spatial transcriptomics technologies that utilise an oligo-dT based transcript capture strategy like most major scRNA-seq workflows ^47^, and so are also compatible with 3’ expressed cellular barcoding methods to enable clonally resolved spatial transcriptomics. Briefly, both BGI Stereo-seq and 10X Genomics Visium, use a grid of coordinate-barcode labelled oligonucleotides printed onto a slide in place of individual cell barcodes linked to gel emulsion beads. Like the capture of lineage barcoded transcripts, this assay setup allows barcode containing reads to be captured by spatial coordinate labelled oligonucleotides and sequenced using high throughput methods, thus linking spatial information with clonal identity.

As a proof of concept of this approach we applied the BGI Stereo-seq strategy to a mouse spleen sample containing SPLINTR barcoded AML cells (hereafter the “spatial” dataset) ^9^. The resulting BGI Stereo-seq data revealed expected splenic morphology (**Figure 5A**) with major Leiden cluster markers revealing the white pulp (Cd74), red pulp (Hba-a1) and interstitial (Marco) regions (**Figure 5B** & **C**). To annotate spatial locations of the mouse spleen with lineage barcode information we applied the *BARtab* “single-cell” workflow to the spatial dataset, specifying the “SAW” aligner using the --pipeline parameter. The Stereo-seq Analysis Workflow (SAW) is a pre-processing pipeline developed by BGI for processing Stereo-seq datasets that links coordinate ID and UMI information to individual transcripts ^48^. Paired spatial coordinate – lineage barcode information was then imported into Scanpy for further analysis. Visualisation of the top 10 most abundant clones (by total number of detected spatial coordinates) in the dataset revealed a restricted pattern of leukaemic clonal outgrowth (**Figure 5D**), with each clone occupying a distinct spatial territory within the spleen section (**Figure 5E**). Given that the representation in individual sections may not reflect the overall clonal representation within a tissue, we compared the frequency of lineage barcodes identified in the spatial dataset to matched population-level and single cell level datasets previously generated from the same mouse spleen ^9^. Of the top 10 most abundant clones in the spatial dataset, all were present in the matched population-level and single-cell level data at similar abundance (**Figure 5F**) supporting the robustness of lineage barcode annotations from the *BARtab* pipeline.

**Figure 5.**
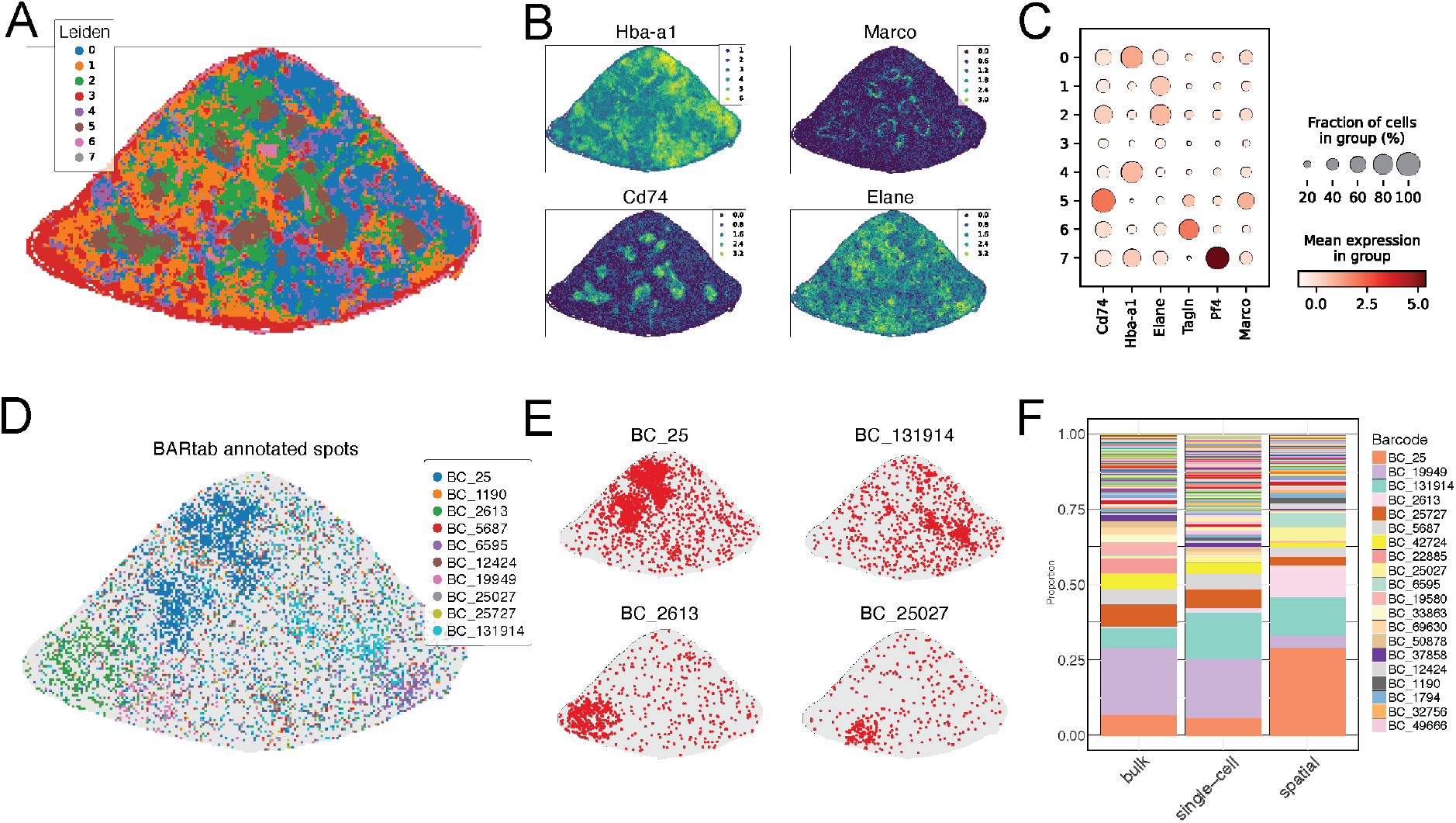
*BARtab* enables clonally resolved spatial transcriptomics. **A)** Spatial section map showing Leiden clustering of the spatial dataset at resolution = 0.7. **B)** Expression pattern of marker genes corresponding to known spleen morphological regions including red pulp (Hba-a1), white pulp (Cd74) and interstitial zone (Marco) and a myeloid marker (Elane). **C)** Dotplot showing average expression of marker genes per Leiden cluster. **D)** Spatial section map showing spatial coordinates annotated to the top 10 most abundant lineage barcodes annotated by *BARtab*. **E)** Spatial section map highlighting spatial coordinates occupied by four of the top 10 most abundant lineage barcodes. **F)** Stacked barplot showing the proportion of clones within the spatial dataset and matched population and single cell level datasets. Most abundant clones are labelled.

## Discussion

Cellular barcoding approaches are widely used in biological research and will increase in utility as new techniques that combine clonal lineage with other cellular modalities such as chromatin accessibility, cell surface protein expression and spatial transcriptomics / epigenomics (e.g. histone modifications and DNA methylation) become more widely available ^49^. These advanced techniques will require the application of robust data analysis tools that can be easily adapted to suit a range of experimental approaches and barcoding systems, and which integrate well with current gold-standard analytical frameworks. The combined workflow provided by *BARtab* and *bartools* solves these data analysis challenges for the field by integrating the pre-processing, quality control and visualisation of cellular barcoding datasets into a readily accessible and flexible workflow that can accommodate many published barcode designs and experimental approaches. Future developments will further enhance the *bartools* and *BARtab* framework to support additional lineage tracing strategies such as CRISPR evolving barcodes ^50,51^, Cre-Lox recombination ^52^, or novel spatial technologies ^14^ to better realise the potential of combining lineage information with the rapidly expanding fields of single-cell and spatial genomics.

## Methods

### Tissue culture

Mouse MLL-AF9 leukaemia cells were generated from the bone marrow of female C56BL6/J mice as previously described ^9,53^ and cultured in RPMI-1640 medium supplemented with mouse IL-3 (10 ng ml^−1^), human IL-6 (10 ng ml^−1^), mouse SCF (50 ng ml^−1^) 20% fetal bovine serum, streptomycin (100 µg ml^−1^), penicillin (100 U ml^−1^) and 2 mM GlutaMAX (Thermo Fisher Scientific) in 5% CO2 at 37 °C. Cell lines were routinely tested for mycoplasma by the Peter MacCallum Genotyping Core Facility and confirmed negative for the duration of the study.

### Dose escalation experiments and analysis

Five hundred thousand mouse MLL-AF9 cells were transduced with the SPLINTR BFP barcode library (Addgene #179776), at a low MOI to ensure single copy integration into cells. BFP positive cells were sorted using a FACS Aria Fusion flow sorter 3 (BD Biosciences) flow cytometer 48 h after transduction. Approximately 5x10^4^ mCherry positive cells were seeded into liquid culture and expanded for seven days. Following expansion, 5x10^5^ cells were harvested as a baseline timepoint zero (T0) sample and processed for population based SPLINTR barcode sequencing as described previously ^9^. To maintain a minimum 20-fold representation of the original theoretical maximum of 5x10^4^ total barcodes per treatment arm, 1x10^6^ cells per replicate and treatment condition were seeded into liquod culture. For the dose escalation arms of the experiment, cultures were supplemented with either 0.1% v/v DMSO, 400nM IBET or 300nM AraC. For the high dose arms, liquid cultures were supplemented with 800nM IBET or 700nM AraC which we had previously determined to represent the equivalent of an IC90 dose in this cell line (data not shown). Drugs were replenished every three days by pelleting the cells at 400 rcf for 5 minutes at 37 degrees Celsius, and replating in fresh medium with drug. For the dose escalation arms, every 7 days, 5×10^5^ cells were replated into fresh medium supplemented with an increased concentration of IBET (TP1 - 400nM, TP-2 - 600nM, TP-3 – 800nM and TP-4 - 1000nM), AraC (TP1 - 300nM, TP-2 - 300nM, TP-3 – 300nM and TP-4 - 500nM) or maintained in 0.1% v/v DMSO. Per timepoint, 1 million cells from each biological replicate were harvested and processed for population based SPLINTR barcode sequencing as described previously.

Raw sequence data from population-based dose-escalation were processed using *BARtab* v1.3 with the following parameters: --mode “single-bulk”, --upconstant “CGATTGACTA”, -- downconstant “TGCTAATGCG”, --alnmismatches 1, --minqual 20, --pctqual 80, --constants “up”, --constantmismatches 0.1, --barcode_length 60. Count files were imported into R v4.2 and further analysed with *bartools* v0.2.5.

### Single cell dataset capture and analysis

5x10^5^ mouse MLL-AF9 cells were transduced with the SPLINTR V0 mCherry barcode library (**Table S2**). Fluorochrome positive cells were isolated using a BD Fusion 5 flow sorter 48 hours after transduction and expanded for seven days in liquid culture as above. Afterward, were processed for single cell transcriptomic capture using the 10x Genomics 3’ V3 Single Cell RNA-seq platform.

Count matrices were generated from demultiplexed scRNA-seq fastq files using the 10x Genomics Cell Ranger (v.3.1.0) count pipeline against the mm10/GRCm38 reference genome. Quality control was performed using Seurat v4 in R v4.2 ^54^. Low-quality cells were removed by filtering out cells that had fewer than 500 genes or 1,000 unique molecular identifiers (UMIs). Cells with greater than 15% mitochondrial RNA content were also removed. For lineage barcode identification and annotation to cells, unmapped reads in BAM file format were extracted from the dataset using Samtools (v1.9) and used as input for *BARtab* v1.3 running in “single-cell” mode with parameters as follows: --mode “single-cell” --pipeline “cellranger”. Downstream analysis was performed with Seurat v4 and *bartools* v0.2.5.

### Spatial transcriptomics capture and analysis

Mouse spleen samples containing SPLINTR barcoded MLL-AF9 leukaemia cells were generated as previously described ^9^ and flash frozen in liquid nitrogen prior to sectioning. BGI Stereo-seq assays were performed using version 1.0 of the Stereo-seq kit and sequenced by BGI genomics group in Shenzhen, China. Section image data was quality controlled with ImageStudio v2.0.1 (BGI). Count matrices were generated from demultiplexed Stereo-seq fastq files using SAW v6.1.0. Reads were aligned to the mm10/GRCm38 reference genome. The Stereo-seq data was manually segmented using the lasso function of StereoMap v2.1.0 (BGI). Stereopy v0.11.0 (BGI) was used to aggregate the count matrix to bin50. Bins with less than 600 UMIs were removed, counts were log transformed and highly variable genes were identified using default parameters in Scanpy v1.9.3 ^34^. Downstream data scaling, PCA and UMAP dimensionality reduction were performed using default parameters. Leiden clustering was performed using resolution = 0.7.

To annotate spatial coordinates with lineage barcodes, unaligned reads resulting from the SAW pipeline were processed with *BARtab* v1.3 using following parameters: upconstant: "TGACCATGTACGATTGACTA", downconstant: "TGCTAATGCGTACTGACTAG", alnmismatches: 2, constantmismatches: 0.2, barcode_length: 60, mode: "single-cell", input_type: "fastq", pipeline: "saw". Barcode counts were aggregated to bin size = 50 and merged with the spatial dataset count matrix based on coordinate ID. For visualization purposes, the barcode with most UMIs per bin was selected.

### Software implementation

*BARtab* and *bartools* are both freely available at https://github.com/DaneVass/bartools and https://github.com/DaneVass/BARtab under MIT and GPL3 licenses respectively. Extensive documentation is available at https://danevass.github.io/bartools. *Bartools* can be installed into R v3.5 or above using instructions that can be found at https://github.com/DaneVass/bartools. *Bartools* integrates with other packages available from the Bioconductor project and utilises functions and object classes from *edgeR* ^35^, *ineq* ^42^ and *vegan* ^41^. Graphical functions from base *R* and *ggplot2* are also utilised within *BARtab* and *bartools*.

*BARtab* can be installed into macOS or UNIX environments compatible with the Nextflow workflow manager v23.04 and later ^23^. *BARtab* depends on common bioinformatic tools including *Samtools* ^27^, *Bowtie* ^25^, *Starcode* ^24^, *FLASh* ^26^, *FastQC* ^29^, *FASTX-toolkit* ^28^, *UMI-tools* ^55^, *MultiQC* ^56^, and GNU parallel ^57^, and requires Python v3.8 or greater. We provide Docker image compatible with Singularity to facilitate pipeline portability across systems. All dependencies required to successfully run the pipeline are also available from the *conda* and *bioconda* projects ^58^. Available parameters for *BARtab* are specified in the documentation (https://github.com/DaneVass/BARtab/blob/main/README.md) which also details approaches for pipeline and software dependency installation via Singularity ^59^, Docker or *conda* environments.

### *BARtab* performance comparison

A dataset consisting of 22 population level barcode sequencing samples from Goyal et al. ^31^ was downloaded from Figshare (https://figshare.com/articles/dataset/FateMap_Paper_datasets_3_Goyal_et_al_2021_Biorxiv_/22806494). The dataset was reanalysed with the published Rewind/TimeMachine code (https://github.com/arjunrajlaboratory/timemachine, commit 9b8c24f) from the same publication using default settings in raw read mode, or with pycashier version v23.1.2 and BARtab v1.3 using settings to mimic these conditions as closely as possible. TimeMachine additionally required the stagger length per sample which was derived from the raw read sequences.

BARtab was run using the following parameters: minqual = 20, pctqual = 80, up_coverage: 10, down_coverage: 10, min_readlength: 40, constants: "both", constantmismatches: 0.1, cluster_distance: 8, cluster_ratio: 5, upconstant: "GACTAAACGCGCTACTTGAT" and downconstant: "ATCCTACTTGTACAGCTCGT".

pycashier was run using the following parameters: quality = 20, unqualified_percent = 20, error = 0.1, length = 100, upstream_adapter = "GACTAAACGCGCTACTTGAT", downstream_adapter = "ATCCTACTTGTACAGCTCGT", ratio = 5, distance = 8, filter_percent = 0 and offset = 8.

For the runtime evaluation, we ran TimeMachine with default settings which clusters barcodes using the barcode read counts. To ensure we ran TimeMachine correctly, we also clustered barcodes on the UMI counts and observed a 100% overlap of identified barcodes with published results. For the comparison of detected barcodes between TimeMachine, BARtab and pycashier, we first aggregated 4 replicates per sample by averaging barcode read counts for BARtab and pycashier. We then filtered barcodes present with at least 0.001% within any sample for all three methods. For the analysis of concordance of barcode quantification between BARtab and TimeMachine, we calculated the spearman and pearson correlation for each sample, considering all barcodes detected in the respective sample by any of the two methods.

## Supplementary Information

Supplementary data are available online.

## Dataset & code availability

The dose escalation, single-cell and spatial datasets have been deposited at the NCBI Gene Expression Omnibus (accession #GSE246611). Code to reproduce the analyses in this manuscript can be found at GitHub (https://github.com/DaneVass/bartools_manuscript_code). The *BARtab* documentation (https://github.com/DaneVass/BARtab/blob/main/README.md) contains information on pipeline installation and execution. The *bartools* documentation (https://danevass.github.io/bartools/) contains further worked examples of cellular barcoding analysis from other previously published datasets, and describes workflows for reference library construction, single cell RNA-seq sample QC, annotation, and analysis.

## Authors contributions

D.V. and H.H. designed and wrote *BARtab* and *bartools* with assistance from E.Y.L. and L.T. D.V., H.H., L.T. and E.Y.L. tested the code and assisted with the development of best practises workflows and analysis vignettes for *BARtab* and *bartools*. K.F. performed experimental work for the dose-escalation and single-cell datasets. D.V., H.H. and L.T. wrote the manuscript with helpful contributions from all other authors. The project was supervised by D.V. and M.A.D. All authors read and approved the manuscript for publication.

## Acknowledgements

The authors would like to acknowledge the support of the Molecular Genomics Core at the Peter MacCallum Cancer Centre, Melbourne Australia for assistance with single-cell sample preparation and sequencing; and BGI Research, Shenzhen, China and BGI Australia for assistance with Stereo-seq library generation, sequencing, and bioinformatics support. The authors also thank the following funders for fellowship, scholarship, and grant support: Leukaemia and Lymphoma Society Career Development Fellowship #3411-22 and Gilead Sciences International Research Scholarship (D.V.). Cancer Council Victoria Sir Edward Dunlop Research Fellowship, NHMRC Investigator Grant #1196749 and Howard Hughes Medical Institute International Research Scholarship #55008729 (M.A.D.); and NHMRC Project Grants #1085015, #1106444 and #1128984 (M.A.D.).

---

**Figure S1.**
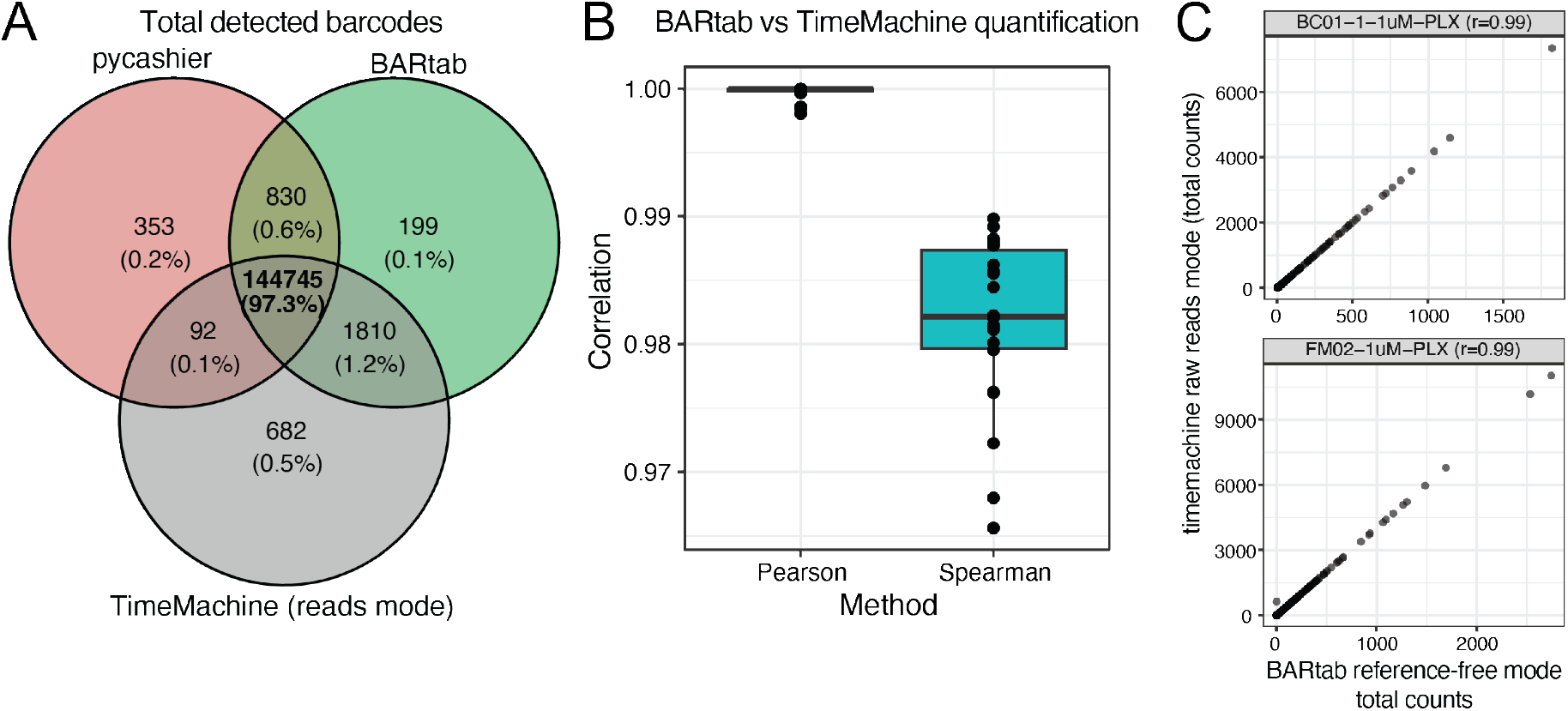
BARtab performance comparison. **A)** Venn diagram of total barcodes detected by BARtab, pycashier and TimeMachine (reads mode) across the 22 samples of the Goyal et al. dataset after filtering barcodes below 0.001% within a sample. B) Pearson and Spearman correlation of BARtab barcode quantification compared to TimeMachine (reads mode). Boxplots indicate mean and interquartile range with whiskers extending 1.5x the IQR. All samples are shown as individual points. C) Exemplar Spearman correlation plots for two samples from the Goyal et al. dataset showing total counts per barcode from BARtab and TimeMachine (reads mode).

**Figure S2.**
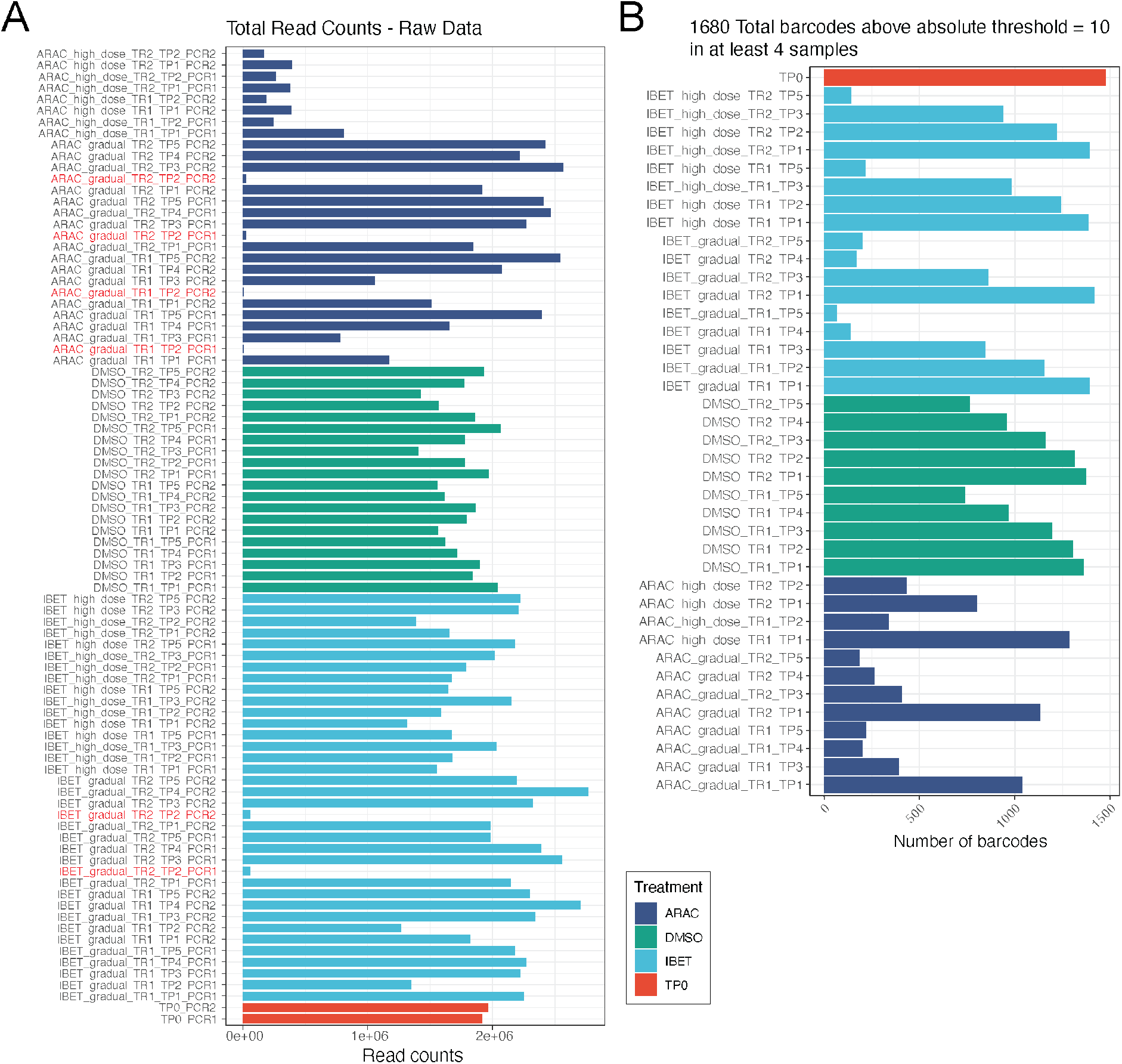
Additional quality control metrics for the dose escalation dataset. **A)** Total number of read counts per sample for the dose escalation dataset. Poor quality samples identified using 5^th^ percentile based outlier detection are highlighted in red. **B)** Total number of lineage barcodes detected per sample post quality control and filtering steps for the dose escalation dataset.

**Figure S3.**
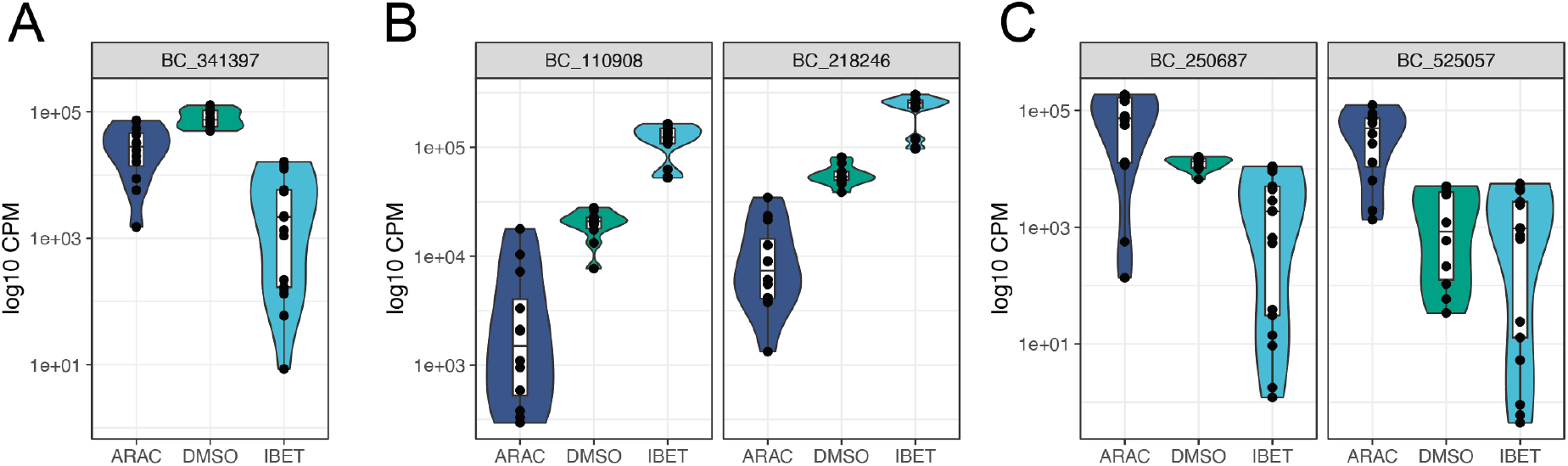
Individual barcode-level analysis. Violin plots of log10 transformed counts per million (CPM) for selected barcodes from the dose escalation dataset predominant in the **(A)** vehicle (DMSO) condition, **(B)** IBET treatment condition and, **(C)** AraC treatment condition. Inset boxplots show the mean and inter-quartile range (IQR). Whiskers extend 1.5x the IQR. Points indicate CPM values for individual samples.

**Figure S4.**
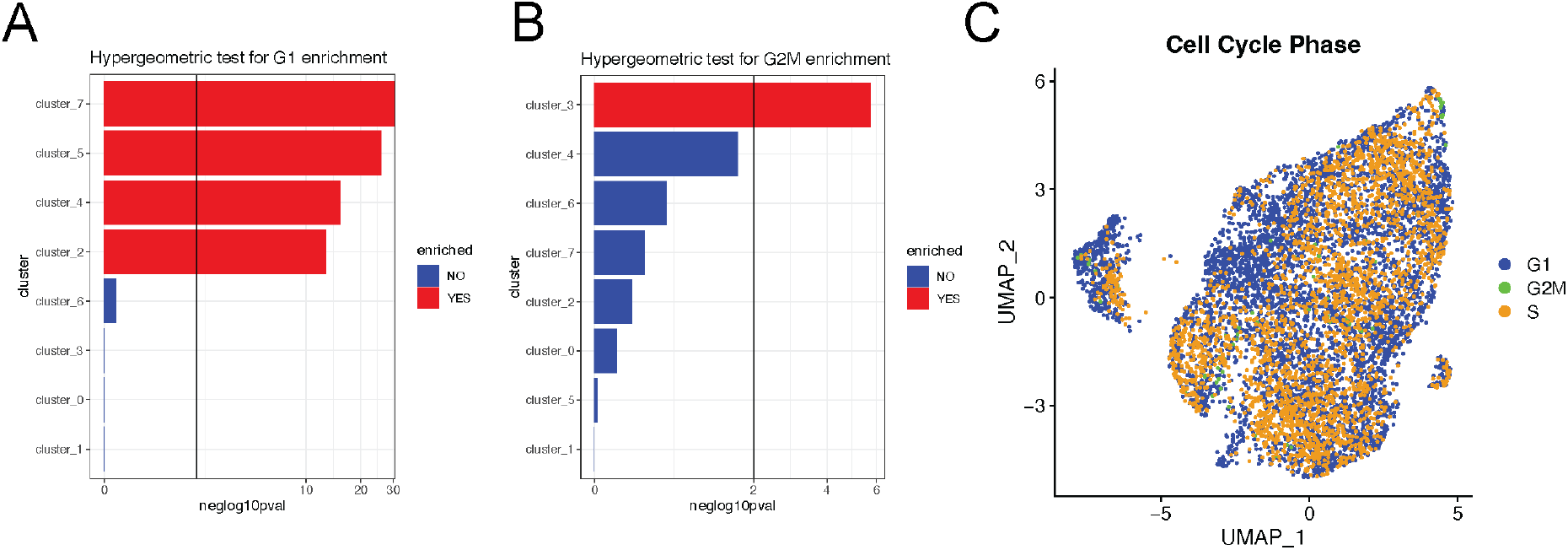
The plotClusterEnrichment function in *bartools* is agnostic to the grouping variable. Hypergeometric testing for enrichment of cell cycle label G1 **(A)** and G2M **(B)** for cells within each Louvain cluster in the single-cell dataset. **C)** UMAP visualisation of the single-cell dataset with cells within each cell cycle phase highlighted.

